# Coordinated Regulation of Renal Glucose Reabsorption and Gluconeogenesis by mTORC2 and Potassium

**DOI:** 10.1101/2024.06.22.600201

**Authors:** John Demko, Bidisha Saha, Enzo Takagi, Anna Mannis, Robert Weber, David Pearce

## Abstract

**Background:** The kidney proximal tubule is uniquely responsible for reabsorption of filtered glucose and gluconeogenesis (GNG). Insulin stimulates glucose transport and suppresses GNG in the proximal tubule, however, the signaling mechanisms and coordinated regulation of these processes remain poorly understood. The kinase complex mTORC2 is critical for regulation of growth, metabolism, solute transport, and electrolyte homeostasis in response to a wide array of inputs. Here we examined its role in the regulation of renal glucose reabsorption and GNG.

**Methods:** Rictor, an essential component of mTORC2, was knocked out using the Pax8-LC1 system to generate inducible tubule specific Rictor knockout (TRKO) mice. These animals were subjected to fasting, refeeding, and variation in dietary K^+^. Metabolic parameters including glucose homeostasis and renal function were assessed in balance cages. Kidneys and livers were also harvested for molecular analysis of gluconeogenic enzymes, mTORC2-regulated targets, and plasma membrane glucose transporters.

**Results:** On a normal chow diet, TRKO mice had marked glycosuria despite indistinguishable blood glucose relative to WT controls. Kidney plasma membrane showed lower SGLT2 and SGLT1 in the fed state, supporting reduced renal glucose reabsorption. Additional metabolic testing provided evidence for renal insulin resistance with elevated fasting insulin, impaired pyruvate tolerance, elevated hemoglobin A1c, and increased renal gluconeogenic enzymes in the fasted and fed states. These effects were correlated with reduced downstream phosphorylation of Akt and the transcription factor FOXO4, identifying a novel role of FOXO4 in the kidney. Interestingly, high dietary K^+^ prevented glycosuria and excessive GNG in TRKO mice, despite persistent reduction in mTORC2 substrate phosphorylation.

**Conclusion:** Renal tubule mTORC2 is critical for coordinated regulation of sodium-glucose cotransport by SGLT2 and SGLT1 as well as renal GNG. Dietary K^+^ promotes glucose reabsorption and suppresses GNG independently of insulin signaling and mTORC2, potentially providing an alternative signaling mechanism in states of insulin resistance.

**SIGNIFICANCE STATEMENT:** The kidney contributes to regulation of blood glucose through reabsorption of filtered glucose and gluconeogenesis. This study shows that mTORC2 and dietary potassium coordinate the regulation of sodium-glucose cotransport and glucose production in the kidney via independent mechanisms. New insights into the regulation of these processes in the kidney offer promising implications for diabetes mellitus management and treatment.

## INTRODUCTION

The proximal tubule (PT) of the kidney contributes to the regulation of blood glucose through reabsorption of filtered glucose and gluconeogenesis (GNG). Sodium-glucose cotransporters 1 and 2 (SGLT1 and SGLT2, respectively) are responsible for reabsorption of filtered glucose across the apical membrane of PT cells, and SGLT2 inhibitors have recently become first-line treatments for type 2 diabetes mellitus (T2DM), proteinuric kidney disease, and heart failure.^1-4^ The liver and the kidney are the only organs capable of GNG in mammals, utilizing substrates such as pyruvate, lactate, and amino acids to synthesize glucose.^5-7^ The kidney is often overlooked but contributes up to 40% of total GNG in healthy individuals.^6, 8^ Both organs exhibit inappropriately increased GNG in the setting of insulin resistance, which is a typical feature of T2DM and the metabolic sydrome.^5-9^ Despite the importance of glucose reabsorption and GNG in human health and disease, relatively little is known about their coordinated regulation in the kidney.

Insulin promotes glucose reabsorption and suppresses renal GNG and in animal models and cultured cells,^10-16^ and humans with T2DM have increased glucose reabsorptive capacity as well as failure to appropriately suppress renal GNG.^17^ Insulin signaling appears to be a common regulatory pathway for glucose reabsorption and renal GNG, however, there is no clear paradigm in the literature due to incomplete characterization of these processes.^11^ Additionally, there may be unique regulatory properties in the kidney because GNG, glucose transport, acid-base homeostasis, and ammoniagenesis are uniquely interconnected within PT cells of the kidney.^18-21^

Glucose is freely filtered across the glomerulus and reabsorbed by the PT, through the same cells that produce de novo glucose.^22^ Apical membrane SGLT2 and SGLT1 mediate glucose reabsorption from the filtrate into PT cells, and the facilitative basolateral glucose transporter 2 (GLUT2) mediates glucose entry into the blood.^22^ The glucose reabsorptive capacity of the PT is increased in humans with type 1 and type 2 diabetes mellitus through unclear mechanisms.^17^ Attempts to explain this phenomenon with diabetic animal models have yielded inconsistent data regarding gene expression and protein abundance of SGLT2, SGLT1, and GLUT2.^11^ In addition, multiple signaling pathways have been implicated in glucose transporter regulation in cell culture without a clear significance *in vivo*.^12-15^ Newer transgenic technology in mice has shown that insulin signaling through the renal insulin receptor, insulin receptor substrates 1 and 2, and Akt2 are important for glucose reabsorption.^10, 16, 23-26^ The precise signaling pathway, importance of mTORC2, and additional regulatory features of glucose reabsorption in the kidney remain active areas of investigation.

Insulin signals through the multiprotein kinase complex mTORC2 to regulate hepatic GNG,^27^ muscle glucose uptake,^28^ and systemic insulin sensitivity.^29, 30^ Insulin activates the insulin receptor on the cell membrane leading to recruitment of mTORC2 and PDK1 to phosphorylate Akt at S473 and T308, respectively.^28, 31^ Both phosphorylation events are necessary for full activity of Akt.^28, 31^ To regulate GNG in the liver, activated Akt phosphorylates and displaces the transcription factor FOXO1 (and possibly other FOXO family members) from the nucleus resulting in decreased gene transcription of the gluconeogenic enzymes phosphoenolpyruvate carboxykinase (PEPCK) and glucose-6-phosphatase (G6Pase).^32, 33^ Insulin signaling through the PT insulin receptor, insulin receptor substrates, and Akt2 are important for suppression of renal GNG.^16, 24-26^ However, the direct role of mTORC2 and downstream signaling changes have not been described in the kidney.

Potassium (K^+^) also affects blood glucose. Hypokalemia worsens glycemic control, which is commonly attributed to reduced insulin secretion from pancreatic beta cells.^34, 35^ However, low serum K^+^ also increases renal GNG and ammoniagenesis in animal models.^18-20^ High dietary K^+^ plays a protective role in human cardiovascular health, and may influence renal glucose production and transport. Large randomized studies have shown that high dietary K^+^ reduces mortality, strokes, and cardiovascular events in humans.^36, 37^ High dietary K^+^ causes natriuresis and reduction in blood pressure, providing some explanation for these benefits, however, epidemiological data also suggests that a high K^+^ diet improves insulin resistance.^35, 37, 38^ High serum K^+^ could suppress renal GNG independently of insulin and mTORC2, making dietary K^+^ beneficial to glycemic control when insulin signaling fails.

Genetically modified mice with inducible renal tubule-specific <u>R</u>ictor <u>k</u>nock<u>o</u>ut (TRKO) have mTORC2 disrupted throughout the tubules by deletion of its essential subunit, Rictor.^39-41^ We have previously shown that TRKO mice are normokalemic on a standard chow diet and hyperkalemic on a high K^+^ diet.^39, 40, 42^ During the course of these studies, we observed that TRKO mice had marked glycosuria on a normal chow diet which resolved on a high K^+^ diet, prompting further characterization of renal glucose transport and GNG. We hypothesized that insulin signaling through mTORC2 is critical for coordinated regulation of renal tubule glucose reabsorption and GNG, and that K^+^ activated an mTORC2-independent bypass pathway to stimulate glucose reabsorption and suppress GNG. We therefore set out to characterize the features underlying coordinated regulation of renal glucose reabsorption and GNG by mTORC2 and K^+^, which might be relevant to the pathogenesis and treatment of T2DM and metabolic syndrome.

## METHODS

### Generation of Inducible Tubule-Specific Rictor Knockout (TRKO) Mice

To study mTORC2 KO throughout the renal tubule, we used C57BL/6 mice with inducible renal tubule-specific Rictor knockout (TRKO) (Pax8-rtTA TetOCre Rictor^flox/flox^).^40^ In this model, Cre recombinase is under the control of a tetracycline operator (TetOCre), and the tetracycline binding protein reverse-tetracycline transactivator (rtTA) is expressed by the renal tubule-specific Pax 8 promotor (Pax8-rtTA).^43^ For Rictor knockout, both copies of exon 11 of the Rictor gene (RPTOR independent companion of mTOR complex 2) are flanked by LoxP sequences (Rictor^flox/flox^).^43^ These two transgenes and the floxed Rictor alleles allow treatment with doxycycline to induce recombination in the renal tubules.^43^

Control mice, referred to as wild-type (WT) mice, were all Rictor^flox/flox^ and lacked either Cre-recombinase or Pax8, a common strategy in the literature.^41^ These mice do not knockout Rictor or reduce mTORC2 activity.^41^ Genotyping was done by PCR-based DNA analysis from tail biopsies as previously described.^40^ In our experiments, TRKO and WT mice between 12-24 weeks of age were treated with two weeks of doxycycline water (2 mg/ml in 2% sucrose drinking water) followed by one week of washout prior to experiments.

### Animal Housing and Sex

All animals were housed in a controlled environment with 12-hour light and 12-hour dark cycles. Mice were raised with free access to water and a standard laboratory diet (#5053, PicoLab Rodent Diet). Approximately equal numbers of male and female mice were used for all experiments, and there were no differences between sexes unless explicitly stated. C57BL/6 mice younger than 24 weeks exhibit no difference in glucose tolerance between sexes.^44^

### Fasting and Refeeding

To elucidate differences in renal glucose transport and GNG, animals were studied during fasting and refeeding in balance cages. Prolonged fasting for 18-hours in mice depletes liver glycogen stores, and maintenance of serum glucose increasingly depends on GNG from the liver and kidneys.^45^ Refeeding was performed by giving mice unrestricted access to food for 4 hours after being denied food for 18 hours. These feeding techniques were used in all experiments to study animals in either the fasted or the fed state.^26, 27^

### Experimental Diets

Prior to all experiments, mice were acclimated for at least one week to a normal chow diet with 0.5% K^+^ and 0.3% Na^+^ (#TD.130041, Teklad diet, Envigo). This diet was also used for refeeding on a normal K^+^ diet. Mice were fed with a diet containing 3% K^+^ and 0.3% Na^+^ for refeeding on a high K^+^ diet (#TD.210203, Teklad diet, Envigo). Both diets have similar amounts of protein, carbohydrates, and fats. They contain an equal ratio of potassium chloride, potassium citrate, and potassium carbonate.

### Balance Cage Studies and Urine Collection

Mice were acclimated to balance cages for two days on a normal chow (0.5% K^+^) diet prior to all experiments. Afterwards, mice were continued on a normal chow diet or switched to a high K^+^ (3% K^+^) diet for all experiments. Mice had unrestricted access to regular drinking water throughout all experiments. Food consumption, water consumption, animal weights, and urine were measured after the acclimation period. For urine collections longer than six hours, urine was collected under water-saturated light mineral oil. For spot urine collections, parafilm was placed underneath the metabolic cages and collected with pipettes at 60-minute intervals as previously described ^25^ All urine was spun at 5000g for two minutes to remove contaminants and stored at -80°C until analysis.

### Blood and Tissue Collection

For repeated glucose measurements, one drop of blood from a tail vein nick in an awake animal was collected and immediately analyzed. For all other blood measurements, whole blood was collected via cardiac puncture under anesthesia and placed into heparinized tubes. The remaining whole blood was centrifuged at 5000g for two minutes, and plasma was stored at - 80°C until further analysis. After blood collection, the kidneys and liver were collected from mice. Connective tissue was dissected off the organs, and blood was washed off with cold PBS. Organs were then snap-frozen in liquid nitrogen and kept at -80°C until further processing. Mice were euthanized by cervical dislocation.

### Blood and Plasma Measurements

Blood glucose was measured immediately from tail vein nicks using a glucometer (Clarity BG1000). Urine glucose was measured in duplicate with the Colorimetric Urine Glucose Assay (STA-680, Cell Biolabs). Electrolytes, hemoglobin, blood urea nitrogen (BUN), and creatinine were measured from heparinized whole blood drawn by cardiac puncture and immediately analyzed using the i-STAT analyzer (Abbott Point of Care). Serum insulin was measured in duplicate from mouse plasma stored at -80°C using the Ultra Sensitive Mouse Insulin ELISA Kit (90080, Crystal Chem). Hemoglobin A1c (HbA1c) was measured in triplicate from whole blood treated with EDTA using the Mouse HbA1c Kit Assay (80310, Crystal Chem).

### Preparation of Kidney and Liver Tissue

Frozen kidneys were homogenized by crushing with a mortar and pestle under liquid nitrogen. Homogenization buffer with phosphatase inhibitors (PhosSTOP 04 906 837 001, Roche) and protease inhibitors (cOmplete Protease Inhibitor Cocktail 11873580001, Roche) were immediately added to the crushed tissue, and homogenization was completed with a Dounce homogenizer and repeated fine needle aspiration. Frozen livers were homogenized in RIPA buffer with protease and phosphatase inhibitors at 30Hz for two minutes in the Tissue Lyser II (Qiagen). Whole tissue lysate was stored at -80°C. For kidneys, an aliquot of whole tissue lysate was used for plasma membrane purification using the Plasma Membrane Purification Kit (ab65400, Abcam) as previously described.^40^ Concentrations of proteins were measured using the Bradford assay and stored at -80°C until western blotting.

### Immunoblotting

Protein abundance was measured using western blot analysis. 40–100µg of whole kidney or 10µg of plasma membrane protein were run using SDS-PAGE and transferred to polyvinylidene difluoride membranes. Incubation with primary antibodies occurred overnight followed by horseradish peroxidase (HRP)-linked secondary antibodies. Band intensities on Hyperfilm were quantified using the Image J software, as previously described.^21^

Primary antibodies included: PEPCK (sc-271029 Santa Cruz), G6Pase (sc-25840 Santa Cruz), SGLT2 (sc-393350 Santa Cruz), GLUT2 (sc-518022 Santa Cruz), phospho-Akt S473 (4060 Cell Signaling Technology (CST)), phospho-Akt T308 (2965 CST), total Akt (9272 CST), phospho-FOXO1 S256 / FOXO4 S193 (9461T CST), phospho-FOXO1 Ser256 (E1F7T CST), total FOXO1 (9454 CST), total FOXO1 (C29H4 CST), total FOXO4 (9472 CST), Pan-cadherin (4068 CST), and ꞵ-actin (4967 CST). SGLT1 antibodies were generously provided by Dr. Koepsell at the University of Würzburg.

### Real-Time PCR

Frozen kidneys and liver were lysed and RNA was extracted using RNeasy mini kits from Qiagen. A cDNA library was prepared using TaqMan reverse transcriptase. Primers for target genes were obtained from previously published sources. cDNA was amplified using the appropriate primers and SYBR Green Master Mix (Thermo Fisher Scientific). Relative mRNA was calculated using the 2^-ΔΔCt^ method after normalizing to actin and using fasted WT mice as the reference as previously described.^27^

Primer sequences included: PEPCK FP 5’-CCCCTTGTCTATGAAGCCCTCA-3’, PEPCK RP 5’-GCCCTTGTGTTCTGCAGCAG-3’, G6Pase FP 5’-TTACCAAGACTCCCAGGACTG-3’, G6Pase RP 5’-GAGCTGTTGCTGTAGTAGTCG-3’, SGLT1 FP 5’-TCTGTAGTGGCAAGGGGAAG-3’, SGLT21 RP 5’-ACAGGGCTTCTGTGTCTTGG-3’, SGLT2 FP 5’-TATTGGTGCAGCGATCAGG-3’, SGLT2 RP 5’-CCCAGCTTTGATGTGAGTCAG-3’, GLUT2 FP 5’-TTTGCAGTAGGCGGAATGG-3’, GLUT2 RP 5’-GCCAACATGGCTTTGATCCTT-3’, ꞵ-actin FP 5’-GGCGGACTGTTACTGAGCTGCG-3’, and ꞵ-actin RP 5’-GCTGTCGCCTTCACCGTTCCA-3’.

### Statistics

GraphPad Prism and Microsoft Excel were used for data analysis. Comparisons between two groups were performed by unpaired t-test. Comparisons of time-dependent data were done by two-way ANOVA with Šidák correction for multiple comparisons. All data are shown as mean ± standard error of the mean (SEM) unless otherwise stated.

### Study Approval

All animal experiments were conducted under the approval of the UCSF Institutional Animal Care and Usage Committee.

## RESULTS

### TRKO Mice have Glycosuria without Hyperglycemia or Metabolic Derangements

Marked glycosuria was consistently observed in TRKO mice on a normal chow diet. Cumulative urine glucose excretion was 19820.37 ±6335.99μg/day in TRKO mice compared to 87.64 ±43.92μg/day in WT mice (p<0.01) (**Table 1**). To determine whether glycosuria was due to hyperglycemia, serial plasma and urine glucose measurements were performed during a 4-hour refeeding experiment and during a 1g/kg intraperitoneal (IP) glucose tolerance test. Blood glucoses were not significantly different at any time point between TRKO and WT mice during refeeding (**Figure 1A**), which was in contrast to hepatic mTORC2 knockout mice which develop hyperglycemia during the same conditions.^27^TRKO mice did have increased urine output, urinary glucose concentration, and cumulative urine glucose excretion compared to WT during refeeding (**Table 1**, **Figure 1B**). Concordant with these results, a glucose tolerance test did not reveal significant differences in serum glucose at any time point between TRKO and WT mice (**Figure 1C**), but cumulative urine glucose excretion was significantly elevated in TRKO mice compared to WT (**Figure 1D**). These data strongly support the conclusion that mice with renal tubule mTORC2 knockout have reduced glucose reabsorptive capacity rather than hyperglycemia as the etiology of glycosuria.

**Figure 1:**
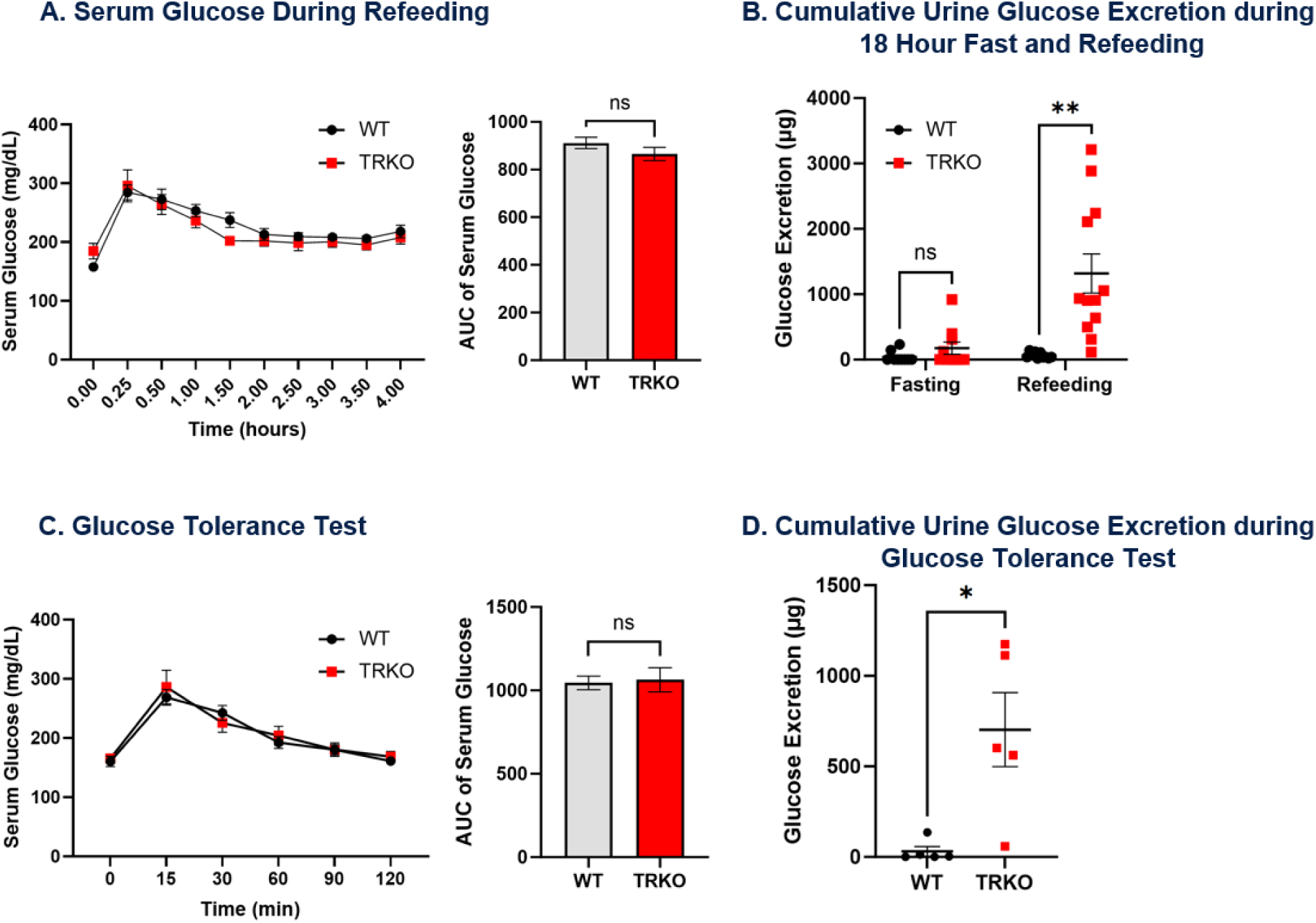
Serum glucose and glycosuria during refeeding and glucose tolerance test in TRKO and WT mice. (A) Serial blood glucose measurements during 4 hours of refeeding with a normal chow diet following 18 hours of fasting (t=0) in TRKO and WT mice. The bar graph to the right shows area under the curve (AUC) of blood glucose during refeeding for each group. (B) Cumulative urine glucose excretion (product of urine glucose concentration and urine volume) after 18 hours of fasting and 4 hours of refeeding with a normal chow diet in TRKO and WT mice. (C) Glucose tolerance test with serial blood glucose measurements after overnight fasting (t=0) followed by intraperitoneal 1g/kg 10% dextrose. The bar graph to the right shows area under the curve (AUC) of blood glucose for each group. (D) Cumulative urine glucose excretion after 2 hour glucose tolerance test. Mice had ad lib access to regular drinking water during all experiments. Values are mean ±SEM; **P <0.01, ns not significant by t-test or 2-way ANOVA with Šidák correction for multiple comparisons in experiments with multiple time points. No significant differences at points without annotations. n=6–10 per group for all experiments.

**Table 1:** Balance, Blood, and Urine Data for WT and TRKO Mice.

|  | 18-Hour Fasting |  | 4-Hour Refeeding |  | 24-Hour Baseline |  |
| --- | --- | --- | --- | --- | --- | --- |
|  | WT | TRKO | WT | TRKO | WT | TRKO |
| **Balance Data** |  |  |  |  |  |  |
| **Weight (g)** | 20.5 ±0.5 | 21.2 ±0.5 | 23.1 ±0.7 | 23.2 ±0.6 | 23.1 ±0.6 | 23.7 ±0.7 |
| **Weight Change (%)** | -11.15 ±0.68 | -10.23 ±0.89 | +11.11 ±1.42 | +9.74 ±1.43 | N/A | N/A |
| **Food Intake (g)** | N/A | N/A | 1.94 ±0.14 | 1.92 ±0.20 | 3.83 ±0.18 | 3.82 ±0.16 |
| **Water Intake (mL)** | 1.70 ±0.32 | 2.41 ±0.30 | 2.43 ±0.17 | 2.62 ±0.21 | 4.56 ±0.28 | 6.24 ±0.39** |
| **Blood** |  |  |  |  |  |  |
| **Na<sup>+</sup> (mmol/L)** | 146 ±1 | 146 ±1 | 146 ±1 | 146 ±1 | 141 ±1 | 141 ±1 |
| **K<sup>+</sup> (mmol/L)** | 4.8 ±0.2 | 4.4 ±0.2 | 4.7 ±0.2 | 4.7 ±0.2 | 5.2 ±0.4 | 5.2 ±0.3 |
| **Cl<sup>-</sup> (mmol/L)** | 115 ±1 | 115 ±1 | 117 ±1 | 118 ±1 | 116 ±1 | 112 ±1 |
| **HCO<sub>3</sub><sup>-</sup> (mmol/L)** | 19 ±1 | 20 ±1 | 20 ±1 | 20 ±1 | 20 ±1 | 23 ±1 |
| **Anion Gap (mmol/L)** | 9 ±2 | 10 ±2 | 9 ±1 | 9 ±1 | 6 ±1 | 6 ±1 |
| **iCal (mmol/L)** | 1.23 ±0.03 | 1.20 ±0.01 | 1.25 ±0.03 | 1.31 ±0.03 | 1.05 ±0.06 | 1.00 ±0.02 |
| **BUN (mg/dL)** | 20 ±2 | 20 ±2 | 31 ±2 | 37 ±2 | 26 ±1 | 28 ±3 |
| **Creatinine (mg/dL)** | <0.2 | <0.2 | <0.2 | <0.2 | <0.2 | <0.2 |
| **Hemoglobin (g/dL)** | 13.1 ±0.6 | 13.7 ±0.3 | 11.5 ±0.3 | 12.1 ±0.2 | 13.1 ±0.2 | 13.2 ±0.4 |
| **Hematocrit (%)** | 38.6 ±1.8 | 40.4 ±0.9 | 33.7 ±0.8 | 35.7 ±0.6 | 38.1 ±0.5 | 39 ±1.3 |
| **Urine** |  |  |  |  |  |  |
| **Urine Output (mL)** | 1.14 ±0.26 | 1.72 ±0.35 | 0.21 ±0.03 | 0.36 ±0.04** | 1.79 ±0.15 | 2.92 ±0.39* |
| **Urine Glucose (mg/dL)** | 8.96<br>±0.03 | 18.00<br>±8.04 | 29.34<br>±4.72 | 385.70<br>±78.66*** | 5.31<br>±2.83 | 592.95<br>±156.62** |
| **Glucose Excretion (ug)** | 41.92<br>±28.64 | 175.18<br>±94.78 | 64.49<br>±14.18 | 1317.01<br>±297.55** | 87.64<br>±43.92 | 19820.37<br>±6335.99** |

All values are mean ±SEM; \*P <0.05, \*\*P <0.01, \*\*\*P<0.001 by t-test. n=6–10 per group for all experiments.

We also assessed basic metabolic parameters in TRKO mice during 18-hour fasting, 4-hour refeeding, and 24-hours on a normal chow diet. All mice appeared active and well-groomed during each experiment. Serum electrolytes, blood urea nitrogen, hemoglobin, and weight were not statistically different between TRKO and WT mice after fasting, refeeding, or 24-hours on a normal chow diet (**Table 1**). Water intake was significantly increased in TRKO mice during the 24-hour collection, and urine output was increased in TRKO mice during refeeding as well as 24-hour assessment (**Table 1**). Food intake did not significantly differ between TRKO and WT mice during refeeding or 24-hour assessment (**Table 1**), which is critical to compare GNG and glucose reabsorption.

### Plasma Membrane SGLT2 and SGLT1 in TRKO Mice are Lower than WT after Refeeding

Glycosuria was only present in TRKO mice in the fed state (**Table 1**, **Figure 1B**). Western blots of kidney plasma membrane fractions showed an approximately 50% lower SGLT2 protein abundance (**Figure 2A**) and approximately 70% lower SGLT1 protein abundance (**Figure 2C**) in TRKO mice relative to WT littermates after refeeding. There was no difference in plasma membrane SGLT2 or SGLT1 protein abundance between TRKO and WT mice after fasting (**Figure 2A and 2C).** SGLT2 mRNA levels were reduced in fasted TRKO mice compared to WT mice (**Figure 2B)**, but there were no significant differences in SGLT2 mRNA levels after refeeding (**Figure 2B**). There were no differences in SGLT1 mRNA levels between TRKO and WT mice after fasting or refeeding (**Figure 2D**). There were no differences in PM GLUT2 protein abundance or mRNA levels between TRKO and WT mice after fasting or refeeding (**Figure 2E and 2F**). These data suggest that glycosuria in TRKO mice is due to relative insufficiency of plasma membrane SGLT2 and SGLT1 in the fed state.

**Figure 2:**
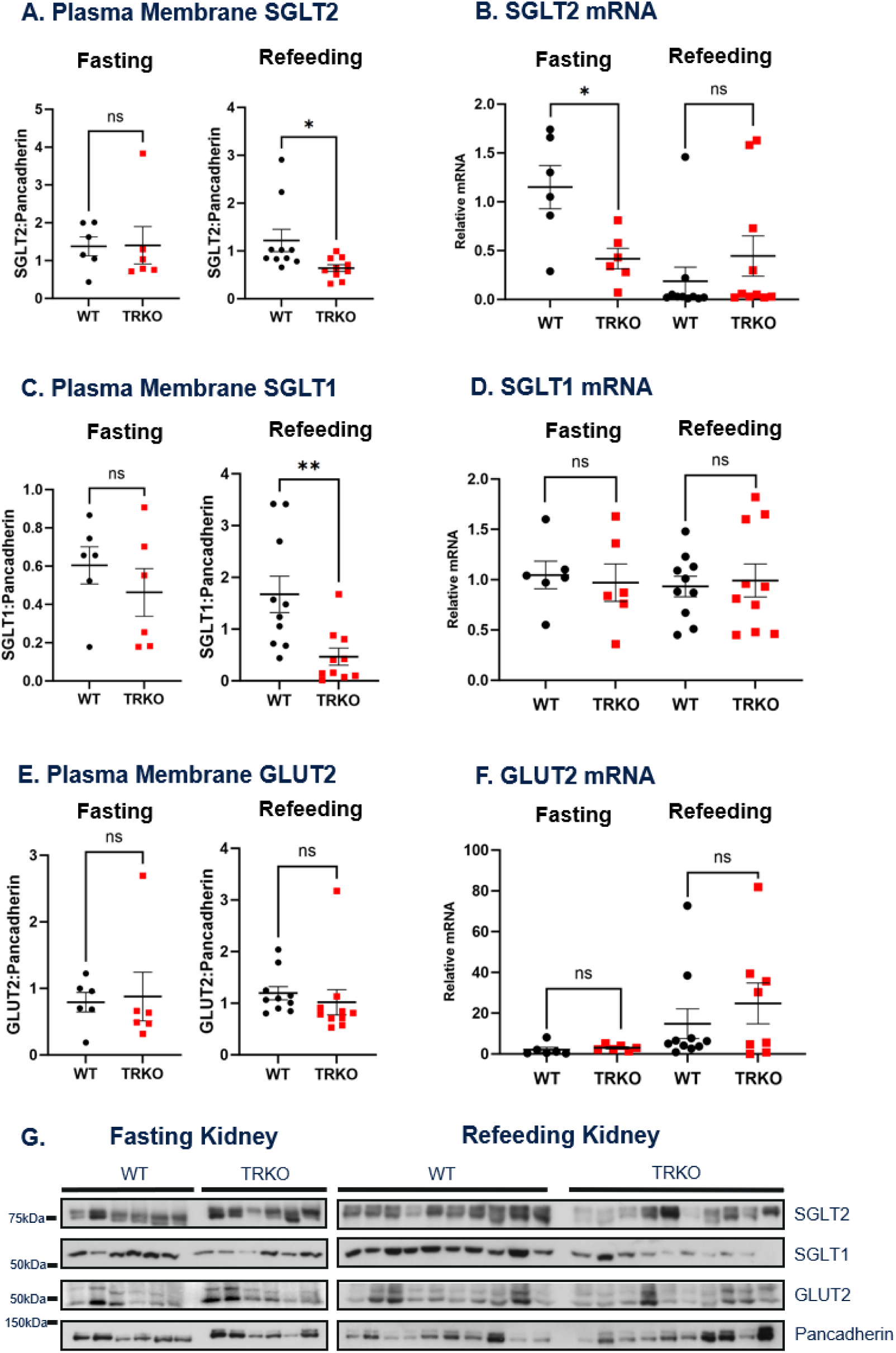
Plasma membrane abundance and expression of renal glucose transporters in TRKO and WT mice after fasting and refeeding. (A) Quantification of PM SGLT2 bands from Western blots and (B) relative SGLT2 mRNA after fasting and refeeding in TRKO and WT mice. (C) Quantification of SGLT1 bands and (D) relative SGLT1 mRNA after fasting and refeeding. (E) Quantification of GLUT2 bands from Western blots and (F) relative GLUT2 mRNA after fasting and refeeding. (G) Original western blots of plasma membrane purified samples from whole kidney homogenate of TRKO and WT mice after 18 hours of fasting (left) and 4 hours of refeeding on normal chow diet (right). PM protein for SGLT2, SGLT1, and GLUT2 were normalized to pancadherin for analysis. Relative mRNA was calculated using the 2^-ΔΔCt^ method after normalizing to actin and using fasted WT mice as the reference. All values are mean ±SEM; *P <0.05, **P <0.01, ns not significant by t-test. n=6–10 per group for all experiments.

### TRKO Mice have Increased Renal Gluconeogenesis

In light of the well-established regulatory role of mTORC2 in hepatic GNG,^27^ it was somewhat surprising that TRKO mice had no obvious abnormality in blood glucose at baseline or during refeeding (**Figures 1A and 1C**). In order to examine TRKO mice for a more subtle abnormality in systemic glucose homeostasis, we fasted the mice overnight and performed additional baseline and tolerance testing, including insulin and pyruvate tolerance tests (**Figure 3**). IP glucose (1g/kg 10%) (**Figure 1C**) and IP regular insulin (0.75U/kg) (**Figure 3A**) did not produce significant differences in blood glucose at any time point between TRKO and WT mice. In comparison, IP pyruvate (2g/kg 10% sodium pyruvate) led to significantly elevated serum glucose at multiple time points TRKO compared to WT mice (**Figure 3B**). Whereas glucose and insulin tolerance tests predominantly reflect acute uptake of glucose by muscle, pyruvate is a substrate for GNG and the pyruvate tolerance test is a well-recognized reflection of GNG.^27^ The most likely site of increased GNG is the PT of the kidney, rather than the liver, due to the selective knockout of mTORC2 in the renal tubules.

**Figure 3:**
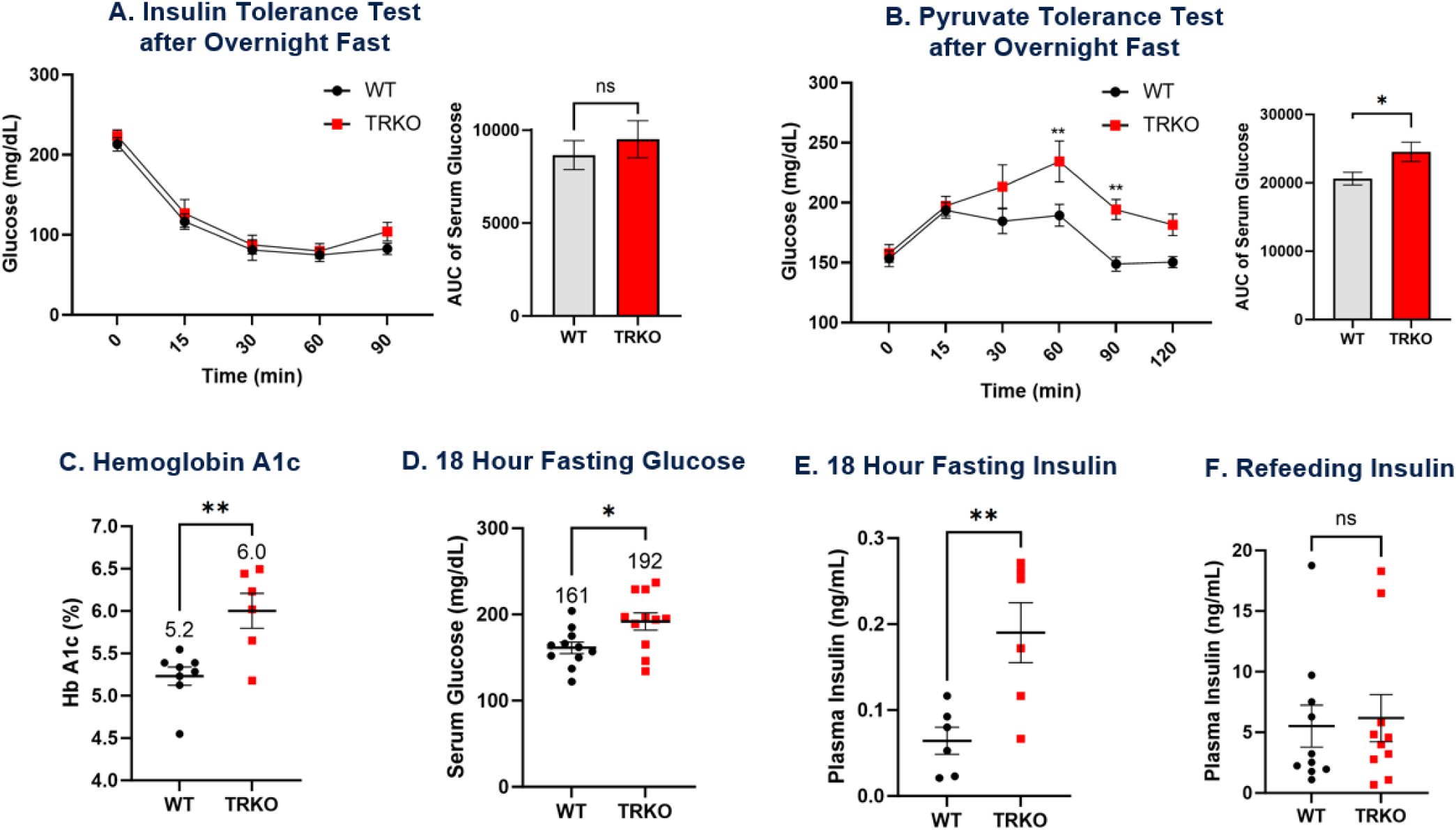
Systemic glucose homeostasis in TRKO and WT mice. Serial blood glucose measurements after overnight fasting (t=0) followed by (A) intraperitoneal 0.75U/kg regular insulin and (B) 2g/kg 10% sodium pyruvate in TRKO and WT mice. The bar graph to the right shows area under the curve (AUC) of blood glucose for each group. (C) Hemoglobin A1c after three months of uninterrupted ad lib access to a normal 0.5% K^+^ diet in TRKO and WT mice. (D) Blood glucose after 18-hour fast in TRKO and WT mice. (E) Serum insulin levels after 18 hours of fasting followed by (F) 4 hours of refeeding with a normal 0.5% K^+^ diet in TRKO and WT mice. All values are mean ±SEM; *P <0.05, **P <0.01 by t-test or 2-way ANOVA with Šidák correction for multiple comparisons in experiments with multiple time points. No significant differences at points without annotations. n=6–10 per group for all experiments.

To assess average serum glucose, we placed TRKO and WT mice on an uninterrupted normal chow diet for three months and measured hemoglobin A1c (Hb A1c). TRKO mice had significantly elevated HbA1c compared to WT animals, with an increase in HbA1c by approximately 0.8% as seen in **Figure 3C**. These data suggest that average blood glucose is higher in TRKO mice and physiologically relevant because hyperglycemia is present with ad lib feeding, not just during experimental conditions.

The relative contribution of renal GNG to total endogenous glucose release increases with prolonged fasting as liver glycogen stores are depleted.^45^ While there was no difference in serum glucose after an overnight fast, TRKO mice compared to WT had significantly elevated fasting serum glucose after a prolonged 18-hour fast (**Figure 3D**). Serum insulin after the 18-hour fast was also significantly elevated in TRKO mice compared to WT (**Figure 3E**), but there was no difference in serum insulin levels after refeeding which reflects a consistent pancreatic response to similar blood glucose levels (**Figure 3F**). In the context of renal tubule specific mTORC2 knockout, hyperglycemia and hyperinsulinemia after prolonged fasting suggest renal insulin resistance in TRKO mice.

### TRKO Mice have Increased Renal Gluconeogenic Enzymes in the Fasted and Fed States

PEPCK and G6Pase gene expression and protein levels are key determinants of GNG. Insulin suppresses GNG by inhibiting FOXO family transcription factors and reducing transcription of the genes encoding these enzymes, such that a fed animal with high glucose and insulin levels will appropriately suppress GNG. Given the evidence for elevated renal GNG in TRKO mice, mRNA and protein abundance of PEPCK and G6Pase were measured after fasting and refeeding in TRKO and WT mice. In the kidney, PEPCK protein levels were significantly elevated in TRKO mice compared to WT after both fasting and refeeding (**Figure 4A**). G6Pase protein levels were only significantly elevated during fasting (**Figure 4C**). In concordance with protein abundance, PEPCK and G6Pase mRNA levels were significantly elevated in TRKO mice after refeeding (**Figure 4B and 4D**). While there was a trend for increased PEPCK mRNA in TRKO mice during fasting, neither PEPCK nor G6Pase mRNA levels were significantly elevated during fasting in TRKO mice (**Figure 4B and 4D**).

**Figure 4:**
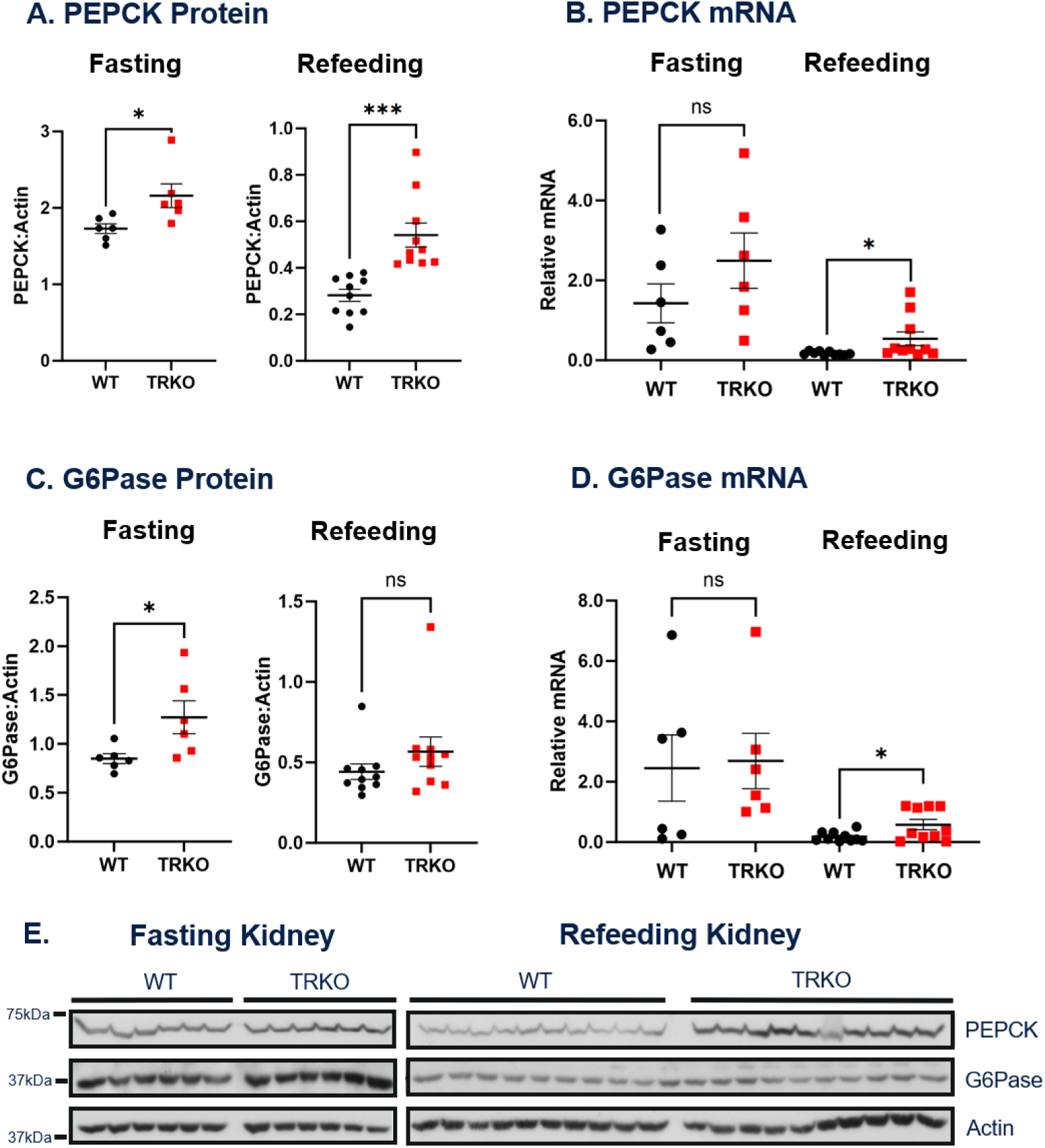
Renal gluconeogenic enzymes in TRKO and WT mice after fasting and refeeding. (A) Quantification of PEPCK bands from Western blots and (B) relative PEPCK mRNA after fasting and refeeding in TRKO and WT mice. (C) Quantification of G6Pase bands and (D) relative G6Pase mRNA after fasting and refeeding. (E) Western blots of whole kidney homogenates for TRKO and WT mice after 18 hours of fasting (left) and 4 hours of refeeding (right) on a normal 0.5% K^+^ diet. PEPCK and G6Pase were normalized to actin for analysis. Relative mRNA was calculated using the 2^-ΔΔCt^ method after normalizing to actin and using fasted WT mice as the reference. All values are mean ±SEM; *P <0.05, **P <0.01, ***P<0.001, ns not significant by t-test. n=6–10 per group for all experiments.

It was unexpected to find a significant increase in PEPCK and G6Pase protein levels after fasting in TRKO mice, but these data are concordant with hyperglycemia seen in TRKO mice after fasting. Increased protein abundance during fasting could be a remnant of increased protein levels in the fed state, increased FOXO-dependent transcription on the fasted state, or unidentified factors increasing protein stability in fasted TRKO mice.

### TRKO Mice have Reduced Akt and FOXO4 Phosphorylation in the Fasted and Fed States

A major mechanism for insulin regulation of GNG, best characterized in liver, is through the kinase cascade that ultimately stimulates mTORC2-dependent phosphorylation of Akt at S473.^28, 31^ For full activation of Akt, it must also be phosphorylated by PDK1 at T308.^28, 31^ In the liver, activated Akt phosphorylates and displaces the gluconeogenic transcription factor FOXO1 from the nucleus, leading to a decrease in PEPCK and G6Pase.^27^ To evaluate this mechanism as well as the importance of mTORC2 in the kidney, phosphorylation of Akt and FOXO1 was measured in TRKO and WT kidneys after fasting and refeeding. Phosphorylation of Akt at S473 was reduced in TRKO animals after both fasting and refeeding; phosphorylation of T308 on Akt was not significantly different after fasting or refeeding in TRKO and WT mice (**Figure 5A and 5B**). Residual phosphorylation of S473 Akt was likely due to cells lacking Cre expression, which have intact mTORC2.

**Figure 5:**
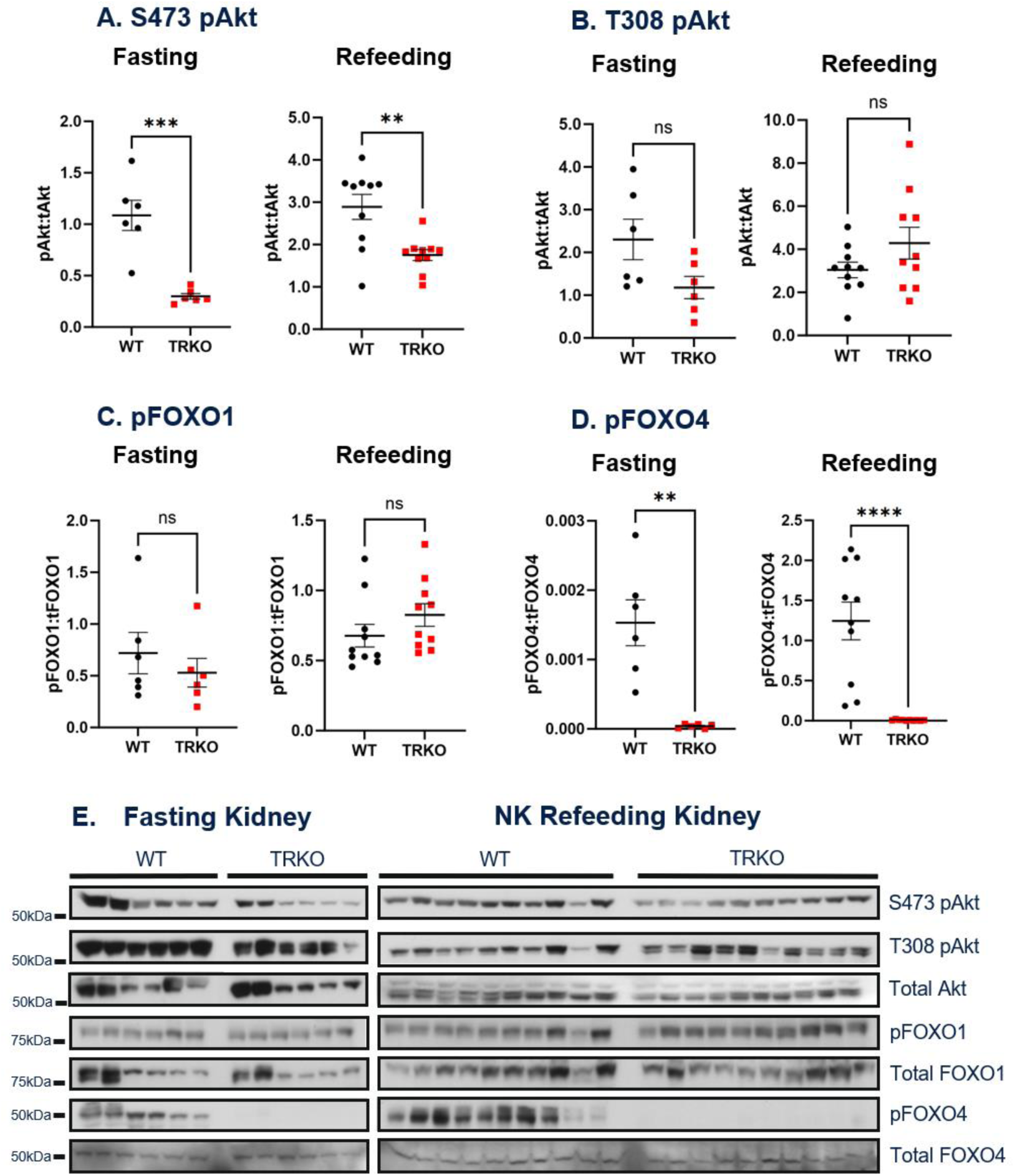
Downstream mTORC2 signaling in TRKO and WT mice. (A-B) Quantification of pAkt bands from Western blots after fasting and refeeding in TRKO and WT mice. (C-D) Quantification of pFOXO1 and pFOXO4 bands from Western blots after fasting and refeeding in TRKO and WT mice. (E) Western blots of whole kidney homogenates for TRKO and WT mice after 18 hours of fasting (left) and 4 hours of refeeding (right) on a normal 0.5% K^+^ diet. S473 pAkt and pT308 Akt were normalized to total Akt for analysis; pFOXO1 and pFOXO4 normalized to total FOXO1 and FOXO4, respectively. All values are mean ±SEM; *P <0.05, **P <0.01, ***P<0.001, ns not significant by t-test. n=6–10 per group for all experiments.

Surprisingly, the phosphorylation of FOXO1 at S256 was not significantly different in TRKO and WT animals after fasting or refeeding (**Figure 5C**), whereas phosphorylation of FOXO1 is significantly reduced in hepatic mTORC2 KO mice.^27^ Instead, we found that phosphorylation of FOXO4 at S193 was dramatically reduced in TRKO mice after both fasting and refeeding (**Figure 5D**). These data suggest that FOXO4, rather than FOXO1, is an mTORC2-dependent transcription factor that regulates glucose homeostasis in the kidney.

### Hepatic Gluconeogenic Enzymes and mTORC2 Signaling in the Fasted and Fed States

We assessed hepatic gluconeogenic enzymes given the liver’s critical role in glucose homeostasis and substantial contribution to GNG. There were no significant differences in hepatic PEPCK or G6Pase protein levels after fasting or refeeding in TRKO and WT mice (**Supplemental Figure 1A and 1C**). However, hepatic PEPCK mRNA was significantly elevated after refeeding in TRKO mice (**Supplemental Figure 1B**). There were no differences in PEPCK mRNA after fasting, and there was no significant difference in G6Pase mRNA after fasting or refeeding (**Supplemental Figure 1B and 1D**). Phosphorylation of Akt at S743 was decreased after fasting in the liver of TRKO mice (**Supplemental Figure 1E**). Conversely, phosphorylation of Akt at T308 was significantly elevated in TRKO livers which is most likely due to hyperinsulinemia in TRKO mice after fasting (**Supplemental Figure 1F**). There were no statistically significant differences in hepatic Akt phosphorylation at S473 or T308 between TRKO and WT mice after refeeding (**Supplemental Figure 1E and 1F**).

Traykova-Brauch *et al* showed that transgenic mice with Pax8-rtTA (such as TRKO mice) express rtTA in periportal hepatocytes.^43^ Therefore, the elevation in PEPCK mRNA after refeeding and decrease in phosphorylation of S473 Akt after fasting are most likely due to partial knockout of mTORC2 in the periportal hepatocytes.^43^ While we cannot completely exclude increased hepatic GNG, the increase in renal gluconeogenic enzymes and unchanged hepatic gluconeogenic enzymes suggest that excess renal GNG is the primary source of hyperglycemia in TRKO mice. Furthermore, these data suggest that the liver is unable to compensate for increased renal GNG, and an interruption in renal insulin signaling alone can cause physiologically significant hyperglycemia.

### Reduced Glycosuria in TRKO Mice on a High K^+^ Diet

Serial serum glucose measurements performed during 4-hour refeeding on a 3% K^+^ diet did not reveal differences between TRKO and WT mice at any time points (**Figure 6A**). In addition, there was no difference in serum insulin after refeeding on a high K^+^ diet (**Figure 6B**). In contrast to the glycosuria seen in TRKO mice on a normal K^+^ diet, the urine glucose concentration and urine glucose excretion were not significantly different between TRKO and WT mice on a high K^+^ diet (**Table 2**, **Figure 6C**). In addition, TRKO mice on 3% K^+^ compared to TRKO mice on a 0.5% K^+^ diet had significantly lower urine glucose concentration (44.32 ±9.91mg/dL vs 385.70 ±78.66mg/dL; p<0.001) and urine glucose excretion in (470.91±115.12μg vs 1317.01 ±297.55μg; p<0.01). These data suggest that serum K^+^ stimulates renal glucose reabsorption, possibly though regulation of glucose transporters.

**Figure 6:**
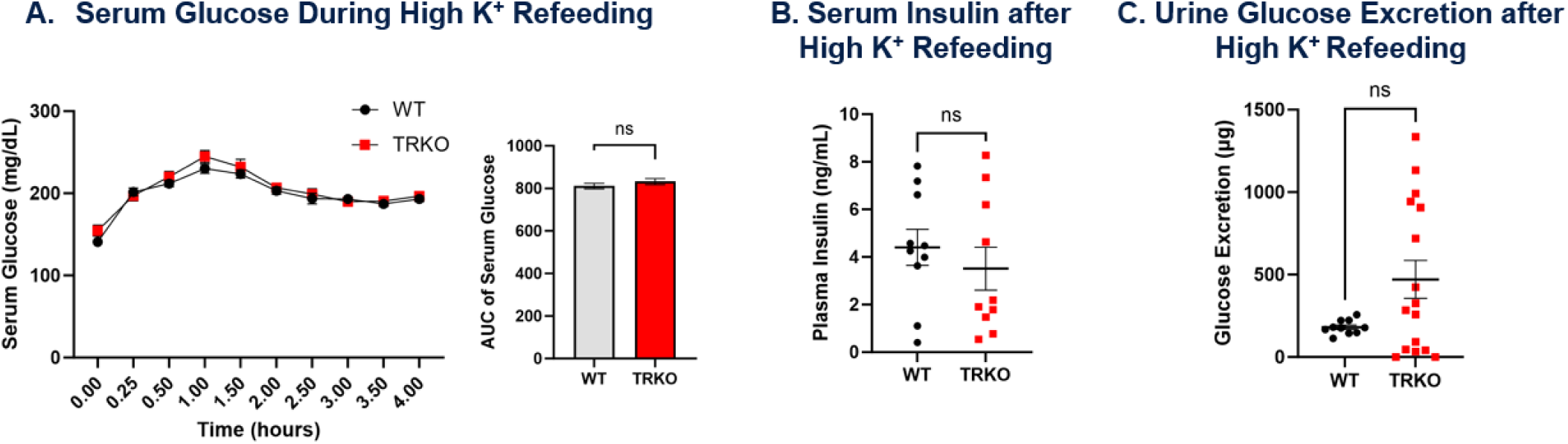
Serum glucose and glycosuria during refeeding on a high K^+^ diet in TRKO and WT mice. (A) Serial blood glucose measurements during 4 hours of refeeding with a high K^+^ diet following 18 hours of fasting (t=0) in TRKO and WT mice. The bar graph to the right shows area under the curve (AUC) of blood glucose during refeeding for each group. (B) Serum insulin levels after 4 hours of refeeding with a high K^+^ diet in TRKO and WT mice. (C) Cumulative urine glucose excretion (product of urine glucose concentration and urine volume) after 18 hours of fasting and 4 hours of refeeding with a high K^+^ Diet in TRKO and WT mice. Mice had ad lib access to regular drinking water during all experiments. Values are mean ±SEM; **P <0.01, ns not significant by t-test or 2-way ANOVA with Šidák correction for multiple comparisons in experiments with multiple time points. No significant differences at points without annotations. n=6–10 per group for all experiments.

**Table 2:** Balance, Blood, and Urine Data for WT and TRKO Mice after High K^+^ Refeeding.

|  | 4-Hour Refeeding on 3% K<sup>+</sup> Diet |  |
| --- | --- | --- |
|  | WT | TRKO |
| **Balance Data** |  |  |
| Weight (g) | 21.27 ±0.81 | 19.58 ±0.94 |
| Weight Change (%) | +5.90 ±0.53 | +3.75 ±0.46** |
| Food Intake (g) | 1.22 ±0.09 | 1.02 ±0.08 |
| Water Intake (mL) | 2.12 ±0.08 | 2.23 ±0.09** |
| **Blood** |  |  |
| Na<sup>+</sup> (mmol/L) | 145 ±1 | 141 ±1*** |
| K<sup>+</sup> (mmol/L) | 6.3 ±0.2 | 7.3 ±0.4* |
| Cl<sup>-</sup> (mmol/L) | 112 ±1 | 111 ±1 |
| HCO<sub>3</sub><sup>-</sup> (mmol/L) | 24 ±1 | 23 ±1 |
| Anion Gap (mmol/L) | 16 ±1 | 15 ±1 |
| iCa (mmol/L) | 1.13 ±0.01 | 1.16 ±0.02 |
| BUN (mg/dL) | 28 ±1 | 33 ±2* |
| Creatinine (mg/dL) | <0.2 | <0.2 |
| Hemoglobin (g/dL) | 13.2 ±0.1 | 14.3 ±0.3** |
| Hematocrit (%) | 38.90 ±0.31 | 42.00 ±0.82** |
| **Urine** |  |  |
| Urine Output (mL) | 0.58 ±0.04 | 1.01 ±0.10* |
| Urine Glucose (mg/dL) | 32.14 ±3.42 | 44.32 ±9.91 |
| Glucose Excretion (ug) | 181.21 ±13.57 | 470.91 ±115.12 |

All values are mean ±SEM; \*P <0.05, \*\*P <0.01, \*\*\*P<0.001 by t-test. n=10 per group for all experiments.

Although all mice appeared active and well-groomed after refeeding on a 3% K^+^ diet, both TRKO and WT mice developed hyperkalemia in this experiment. TRKO mice had significantly higher serum K^+^ and evidence of volume depletion compared to WT animals (**Table 2**). These data are consistent with impaired K^+^ excretion in the distal nephron due to mTORC2 knockout throughout the renal tubules, as previously described.^39, 40, 42^ Food intake remained comparable between TRKO and WT mice (**Table 2**), allowing for comparison of glucose homeostasis, glucose reabsorption, and GNG.

### Plasma Membrane SGLT2 and SGLT1 are Similar in TRKO and WT Mice on a High K^+^ Diet

In contrast to refeeding on a normal K^+^ diet, there were no differences in plasma membrane abundance of SGLT2, SGLT1, or GLUT2 between TRKO and WT mice after refeeding with a 3% K^+^ diet (**Figure 7A, 7C, and 7E**). There were also no differences in mRNA levels of SGLT2 or SGLT1 during this experiment (**Figure 7B and 7D**). Unexpectedly, GLUT2 mRNA levels were significantly lower in TRKO compared to WT mice after 3% K^+^ refeeding (**Figure 7F**). Whereas TRKO mice on a normal K^+^ diet have significantly reduced plasma membrane SGLT2 and SGLT1, plasma membrane SGLT2 and SGLT1 in TRKO mice on a high K^+^ diet is indistinguishable from WT, which is a likely reason for the marked improvement in glycosuria. In addition, these data suggest that K^+^ is able to restore sodium-glucose cotransport independently of mTORC2.

**Figure 7:**
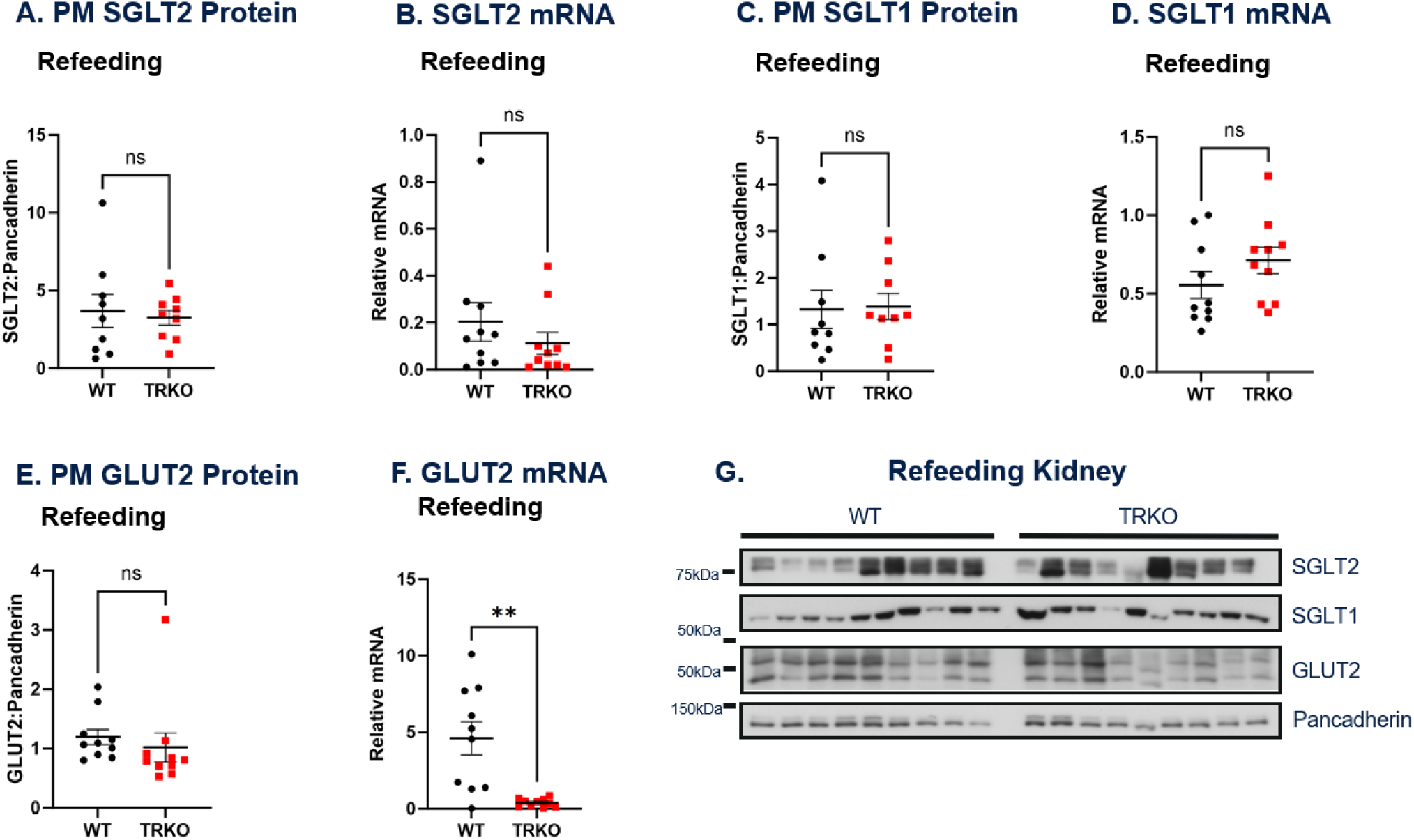
Plasma membrane abundance and expression of renal glucose transporters in TRKO and WT mice after refeeding on a high K^+^ diet. (A) Quantification of PM SGLT2 bands from Western blots and (B) relative SGLT2 mRNA after refeeding in TRKO and WT mice. (C) Quantification of SGLT1 bands and (D) relative SGLT1 mRNA after refeeding. (E) Quantification of GLUT2 bands from Western blots and (F) relative GLUT2 mRNA after refeeding. (G) Western blots of plasma membrane (PM) purified samples from whole kidney homogenate of TRKO and WT mice after 18 hours of fasting and 4 hours of refeeding on a high 3% K^+^ diet. PM protein for SGLT2, SGLT1, and GLUT2 were normalized to pancadherin for analysis. Relative mRNA was calculated using the 2^-ΔΔCt^ method after normalizing to actin and using fasted WT mice as the reference. All values are mean ±SEM; *P <0.05, **P <0.01, ns not significant by t-test. n=6–10 per group for all experiments.

### TRKO Mice have Suppressed Renal GNG on a High K^+^ Diet

In contrast to refeeding on a normal K^+^ diet, PEPCK protein abundance and mRNA levels in the kidney were significantly lower in TRKO animals compared to WT after high K^+^ refeeding (**Figure 8A and 8B**). G6Pase protein and mRNA levels were also significantly reduced in TRKO mice on a high K^+^ diet (**Figure 8C and 8D**). The suppression of gluconeogenic enzyme mRNA levels in these experiments suggest regulation at the level of transcription or transcript stability. The data does not exclude additional regulatory features at the protein level. TRKO and WT mice did not have a statistically significant differences in food intake or serum insulin levels during this experiment, so it is unlikely that either of these factors explain the significant suppression of gluconeogenic enzymes in TRKO mice (**Table 2**, **Figure 6B**). TRKO mice had more severe hyperkalemia compared to WT littermates, most likely due to impaired K^+^ excretion in the distal nephron (**Table 2**).^39, 40, 42^ It is possible that the higher serum K^+^ concentration was responsible for suppression of renal gluconeogenic enzymes in TRKO mice, though additional experiments are needed to elucidate the relationship between serum K^+^ concentration and regulation of renal GNG.

**Figure 8:**
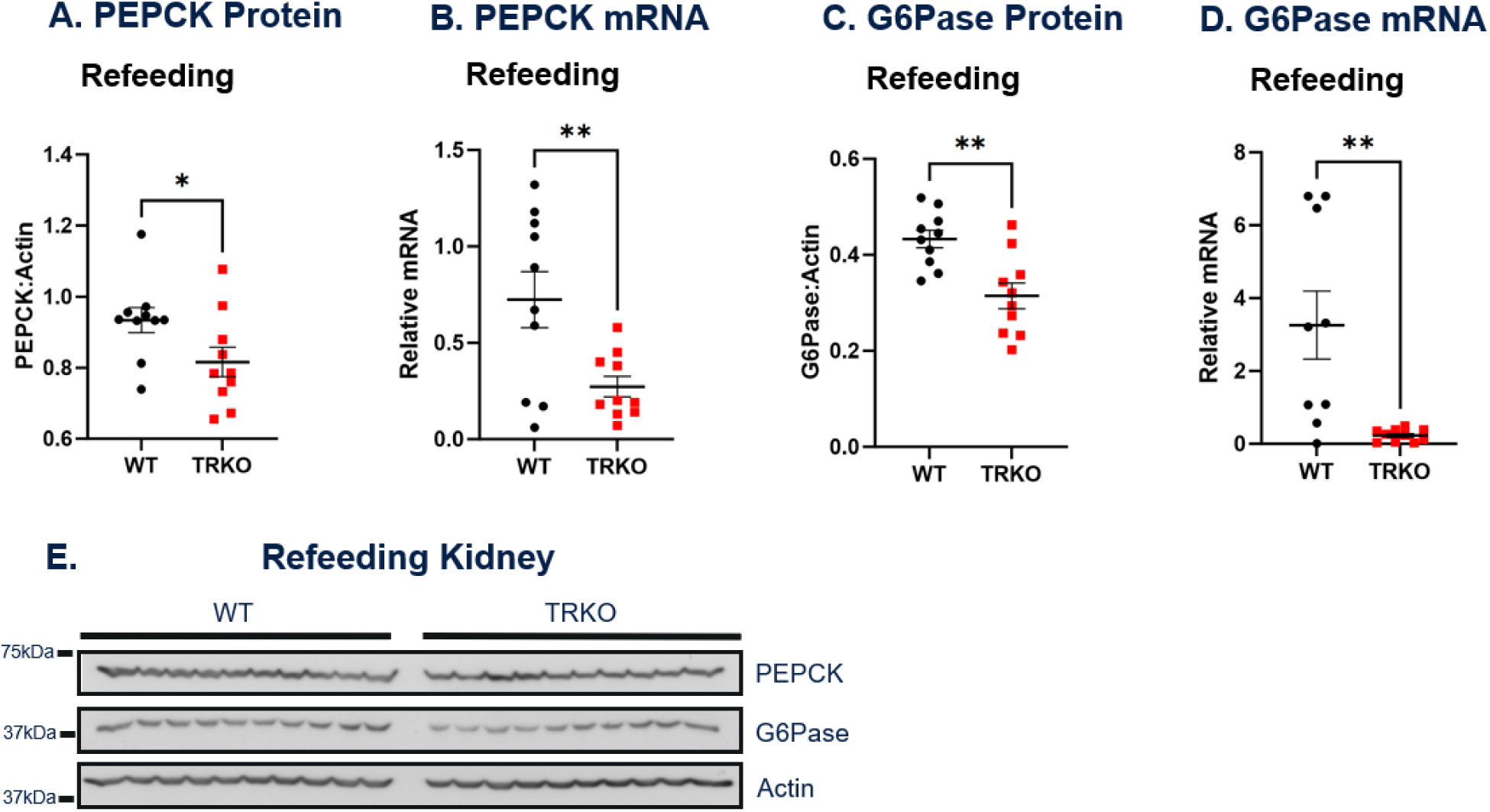
Renal gluconeogenic enzymes in TRKO and WT mice after refeeding on a high K+ diet. (A) Quantification of PEPCK bands from Western blots and (B) relative PEPCK mRNA after refeeding in TRKO and WT mice. (C) Quantification of G6Pase bands and (D) relative G6Pase mRNA after refeeding. (E) Western blots of whole kidney homogenates for TRKO and WT mice after 18 hours of fasting and 4 hours of refeeding on a high 3% K^+^ diet. PEPCK and G6Pase were normalized to actin for analysis. Relative mRNA was calculated using the 2^-ΔΔCt^ method after normalizing to actin and using fasted WT mice as the reference. All values are mean ±SEM; *P <0.05, **P <0.01, ***P<0.001, ns not significant by t-test. n=6–10 per group for all experiments.

### TRKO Mice have Reduced Akt and FOXO4 Phosphorylation after High K^+^ Refeeding

There was a reduction in phosphorylation of Akt at S473 in TRKO animals on a high K^+^ diet, which was similar in magnitude to the reduced phosphorylation at this site in TRKO animals on a normal K^+^ diet (**Figure 9A**). There was no significant difference in phosphorylation of Akt at T308 (**Figure 9B**). Therefore, the suppression of PEPCK and G6Pase on a high K^+^ diet appears to be independent of insulin and mTORC2 signaling.

**Figure 9:**
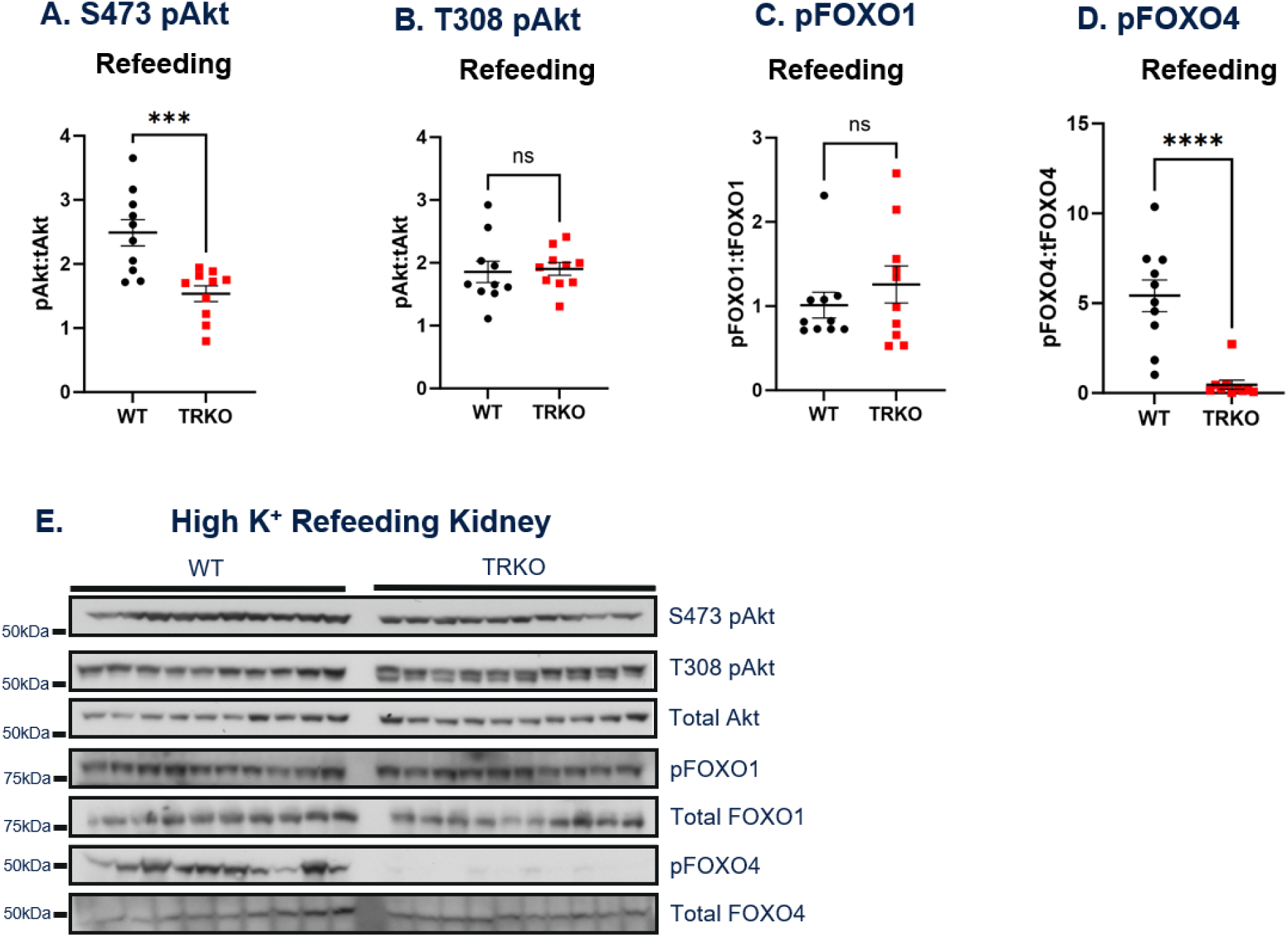
Downstream mTORC2 signaling in TRKO and WT mice on a high K^+^ diet. (A-B) Quantification of pAkt bands from Western blots after refeeding in TRKO and WT mice. (C-D) Quantification of pFOXO1 and pFOXO4 bands from Western blots after refeeding in TRKO and WT mice. (E) Western blots of whole kidney homogenates for TRKO and WT mice after 4 hours of refeeding on a high 3% K^+^ diet. S473 pAkt and pT308 Akt were normalized to total Akt for analysis; pFOXO1 and pFOXO4 normalized to total FOXO1 and FOXO4, respectively. All values are mean ±SEM; ***P<0.001, ****P<0.0001, ns not significant.

There was also no difference in phosphorylation of FOXO1 at S256 between TRKO and WT mice after high K^+^ refeeding (**Figure 9C**). However, phosphorylation of FOXO4 at S193 was markedly decreased in TRKO mice after high K^+^ refeeding (**Figure 9D**). The phosphorylation pattern of FOXO1 and FOXO4 was similar in TRKO mice after both normal and high K^+^ refeeding. These data suggest that high K^+^ acts independently of FOXO1 and FOXO4 to regulate renal glucose homeostasis, but more rigorous experiments are needed to fully elucidate the relationship between K^+^ and FOXO4 in the PT.

### Hepatic Gluconeogenic Enzymes and mTORC2 Signaling after High K^+^ Refeeding

To assess the sensitivity of hepatic GNG to dietary K^+^, gluconeogenic enzymes and the insulin signaling pathway were assessed after high K^+^ refeeding. There were no differences in hepatic PEPCK or G6Pase protein levels or mRNA levels between TRKO and WT mice (**Supplemental Figure 2A-D**). There was a decrease in hepatic S473 pAkt in TRKO mice which was also seen after normal K^+^ refeeding (**Supplemental Figure 2E**). There were no differences in hepatic T308 pAkt between TRKO and WT mice (**Supplemental Figure 2F**). These data are consistent with a prior study that showed hepatic GNG is not sensitive to changes in serum pH or K^+^.^20^

## Discussion

Although it has become increasingly clear that renal GNG and glucose reabsorption play an important role in systemic glucose homeostasis, these processes and in particular their coordinated regulation remain incompletely understood. From an evolutionary perspective, it is logical that animals would suppress renal GNG and increase glucose reabsorptive capacity after a meal. This coordinated regulation prevents unnecessary (metabolically costly) glucose production and minimizes urinary glucose loss in this energy rich state. Recently, mTORC2 has become a focal point for studying insulin signaling and insulin resistance in liver, pancreas, muscle, hypothalamus, and adipose tissue.^27-30^ Using tubule-specific mTORC2 knockout models in mice, we have now demonstrated the essential role of renal mTORC2 in the coordinated regulation of GNG and glucose transport and elucidated its impact on systemic glucose homeostasis.

In TRKO mice lacking mTORC2 throughout the renal tubules (via deletion of the essential subunit Rictor), glycosuria was consistently observed in the fed state despite a lack of significant differences in serum glucose between TRKO and WT mice (**Figure 1**). Since there was no evidence that glomerular filtration rates were different, this indicates that the filtered load of glucose was comparable between TRKO and WT mice and strongly suggests that the glycosuria is due to a tubular transport defect. In further support of this conclusion, plasma membrane expression of SGLT2 and SGLT1 were significantly lower in TRKO compared to WT mice after refeeding (**Figure 2**).

Several studies have shown that insulin increases total protein abundance of SGLT2 and SGLT1 in animal models and cultured cells, however, regulation of PM abundance in response to fed and fasted states is poorly characterized.^14, 15, 46^ Phosphorylation of SGLT2 and SGLT1 are potential regulatory steps to maintain PM protein expression. Specifically, insulin-induced phosphrylation of SGLT2 at Ser-624 has been shown to increase half-life at the PM *in vitro*.^22, 47, 48^ Regulation of SGLT1 at the PM is similarly increased by Akt2, PKA, and PKC *in vitro*, although a consensus phosphorylation site has not been identified.^15, 22, 23^ Newer transgenic technology in mice show that insulin signaling through the renal insulin receptor and Akt2 is important for glucose reabsorption *in vivo*, but the role of mTORC2 and regulation of PM sodium-glucose cotransporters has yet to be defined.^10, 23^ The lower abundance of PM SGLT2 and SGLT1 in TRKO mice after refeeding is most likely due to disruption of downstream insulin signaling via mTORC2 deletion, although it is unclear if mTORC2, Akt2, or another downstream kinase is directly responsible for phosphorylation of these cotransporters. Additional experiments are needed to fully elucidate the signaling cascade and mechanism(s) which regulate SGLT2 and SGLT1 in the kidney.

Hyperglycemia was not observed in TRKO mice during refeeding, glucose tolerance testing, or insulin tolerance testing (**Figure 1A, 1C, and 3A**). Intact muscle uptake of glucose and loss of glucose in the urine is the most likely explanation for normal serum glucose during these experiments. However, we found that mild hyperglycemia and hyperinsulinemia were present after prolonged fasting in TRKO mice (**Figure 3**). The average HbA1c was also 0.8% higher in TRKO mice compared to WT, indicating an increased average blood glucose in TRKO mice (**Figure 3C**). To assess the etiology of hyperglycemia, we performed pyruvate tolerance testing and found that TRKO mice developed hyperglycemia which is consistent with increased GNG (**Figure 3B**). Furthermore, we found that gluconeogenic proteins in the kidney were higher during fasting and refeeding in TRKO mice compared to WT mice (**Figure 4**). Together these data offer compelling evidence that renal tubule mTORC2 knockout leads to hyperglycemia via increased GNG, and increased transcription of these enzymes due to mTORC2 KO is the most likely mechanism, as proposed in **Figure 10**.

**Figure 10:**
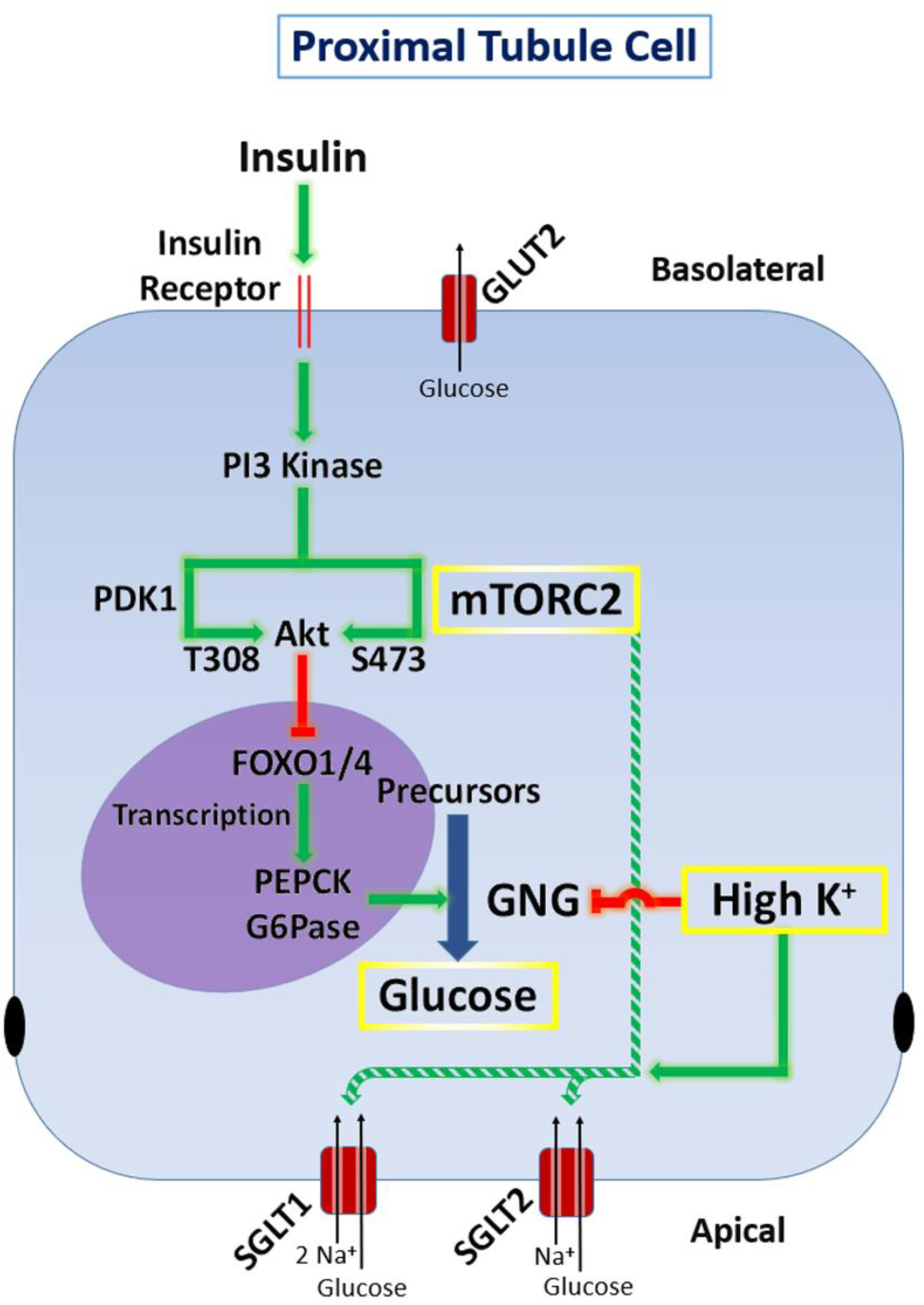
Proposed regulation of gluconeogenesis and glucose reabsorption by mTORC2 and K^+^ in the proximal tubule. Insulin leads to Akt phosphorylation by mTORC2 and PDK1 at distinct sites, both are required for full activity. Akt phosphorylates FOXO1 and displaces it from the nucleus leading to decreased transcription of PEPCK and G6Pase and ultimately decreased GNG. Alkaline pH and high K also inhibit GNG. The pathway for GNG is simplified for clarity. The dashed green line shows that mTORC2 increases plasma membrane SGLT2, but the precise mechanism is unknown.

When we examined the gluconeogenic transcription factor FOXO1, it was unexpected to find no differences in Akt-dependent phosphorylation between TRKO and WT mice after fasting and refeeding (**Figures 5 and 9**). FOXO1 is a well-established substrate of Akt in the liver, and reduced mTORC2-dependent phosphorylation of Akt at S473 is sufficient to decrease Akt activity.^27, 28, 31, 40, 49-52^ Prior studies have also suggested that disruption of insulin signaling in the mouse kidney reduces FOXO1 phosphorylation at S256.^25, 26^ However, these studies did not explicitly differentiate proteins by molecular weight on western blots, and the antibody (CST #9461) used in these studies cross-reacts with FOXO4 at S193 according to the manufacturer and peptide sequence.^53^ We found a dramatic reduction in pFOXO4 in TRKO mice after all fasting and feeding conditions using the same antibody (CST #9461), differentiating pFOXO1 and pFOXO4 by molecular weight (**Figures 5 and 9**). While relatively little is known about the regulatory functions of FOXO4, it has been reported to increase gluconeogenic gene transcription in the liver.^50, 54^ To our knowledge, these data suggest a novel and mTORC2-dependent role of FOXO4 in regulation of renal gluconeogenic gene expression, and the kidney differs from the liver which involves FOXO1 in this pathway. Furthermore, FOXO4 may play a unique role in coordinating the regulation of GNG and sodium-glucose cotransporters in the kidney. Additional experiments are required to confirm and further explore the regulatory role of FOXO4 in the kidney.

While investigating the role of mTORC2 and distal nephron K^+^ excretion,^39, 40, 42^ we observed that glycosuria dramatically improved on a high K^+^ diet in TRKO mice. Given the well-established benefits of high dietary K^+^ on human cardiovascular health and mortality, we further investigated the role of dietary K^+^ on renal glucose transport and GNG in the context of mTORC2 knockout.^36, 37^ We hypothesized that K^+^ bypassed mTORC2 to stimulate glucose reabsorption and suppress GNG through an insulin-independent signaling mechanism. It remains mechanistically unclear how stimulation of sodium-glucose cotransport by K^+^ would lead to human health benefits in light of the robust clinical benefits seen with SGLT2 inhibitors.^1- 4^ However, a signaling pathway for suppression of GNG that remained intact in states of insulin resistance would offer a new approach for the treatment of T2DM, because insulin fails to completely suppress GNG in the kidney and liver in in T2DM.^8^

While TRKO mice developed glycosuria while refeeding with normal chow, urine glucose excretion was not significantly different between TRKO and WT mice after refeeding with a high K^+^ diet (**Table 2**, **Figure 6C**). It is likely that glycosuria resolved because there was normalization of PM SGLT2 and SGLT1 on this diet (**Figure 6**). In contrast, TRKO animals refed on a normal K^+^ diet had glycosuria and lower PM SGLT2 and SGLT1 (**Figure 2**). The persistently reduced phosphorylation of Akt at S473 and FOXO4 at S193 in TRKO mice after high K^+^ refeeding (**Figure 9A**) suggests that K^+^ acts through a bypass pathway that does not involve mTORC2, Akt2, of FOXO4.

Regarding GNG, refeeding on a high K^+^ diet resulted in suppressed PEPCK and G6PAse protein levels as well as mRNA levels in TRKO compared to WT mice. The opposite result was seen after refeeding on a normal K^+^ diet, where gluconeogenic enzymes were increased in TRKO mice. Furthermore, there were no differences in food intake, serum glucose, or serum insulin between TRKO and WT mice after refeeding on a high K^+^ diet (**Table 2**, **Figure 6**). Phosphorylation of Akt at S473 and FOXO4 at S193 were similarly reduced in TRKO mice regardless of dietary K^+^ content (**Figure 5 and 9**). In summary, these data suggest that high serum K^+^ can suppress renal GNG and promote sodium-glucose cotransport, and the mechanism is independent of insulin and mTORC2 signaling.

There is a paucity of data regarding the relationship between K^+^ and glucose homeostasis in humans and animals, and most mechanistic research has focused on the relationship between low serum K^+^ and glucose homeostasis. Hypokalemia is thought to worsen glycemic control by reducing insulin secretion from pancreatic beta cells.^34, 35^ In addition, low serum K^+^ also increases renal GNG and ammoniagenesis in animal models.^18-20^ Cell hyperpolarization due to low serum K^+^ leads to increased basolateral exit of bicarbonate via the electrogenic sodium bicarbonate cotransporter 1 (NBCe1), and therefore low serum K^+^ decreases intracellular pH in the PT.^21^ Acidic intracellular pH increases ammoniagenesis and GNG in the kidney through unclear mechanisms.^18-20^ Notably, the increase in renal GNG is mediated by an increase in gluconeogenic enzyme activity and availability, rather than substrate availability.^18^

Additional experiments are needed to elucidate the mechanism by which K^+^ regulates renal glucose transport and GNG. There are no reports of K^+^ influencing renal glucose transport directly, but regulation at the protein level is likely because there were no differences in SGLT2 or SGLT1 mRNA between TRKO and WT mice during refeeding on a normal or high K^+^ diet. Regulation of PEPCK transcript stability by pH has previously been described.^55^ Alkaline pH within the PT from high extracellular K^+^ may stabilize the transcripts for PEPCK and G6Pase, and this hypothesis would be consistent with our findings (**Figure 7**). There were no differences observed in hepatic PEPCK or G6Pase after refeeding on a high K^+^ diet, which is also consistent with prior reports.^20^ Additional experiments with high dietary K^+^ was not feasible in our model, because prolonged exposure to a high K^+^ diet leads to metabolic derangements and volume depletion in TRKO mice due to mTORC2 knockout in the distal nephron.^39, 40, 42^ Therefore, future experiments with mTORC2 KO isolated to the proximal tubule will be needed to fully understand the effects of dietary K^+^ on systemic glucose homeostasis.

The current evidence shows the direct role of mTORC2 in the coordinated regulation of renal GNG and sodium-coupled glucose reabsorption via SGLT2 and SGLT1, as proposed in **Figure 10**. We have furthermore exposed a striking difference between renal and hepatic signaling mechanisms by identifying FOXO4 as the central mediator of mTORC2 signaling, distinct from liver which uses FOXO1. It seems likely that this difference is in support of the coordinated regulation of GNG and sodium-glucose cotransport in the kidney. In addition, we found a novel role of high dietary K^+^ in the regulation of renal GNG and glucose transport, which appears to be independent of insulin and mTORC2. These data have promising implications for management of hyperglycemia via mTORC2-independent suppression of renal GNG by K^+^. Future work will build on this evidence to elucidate the regulatory mechanisms of renal GNG and glucose reabsorption by K^+^ as well as the potential benefits of high dietary K^+^ in the setting of insulin resistance.

## Author contributions

JD and ET conducted the experiments. JD and DP designed the study. JD, BS, and DP analyzed data. JD, RW, and DP drafted the manuscript.

## Acknowledgements

We thank Prof. Hermann Koepsell (University of Wuerzburg, Germany) for providing the SGLT1 antibody. We are grateful to Dr. Carolyn Ecelbarger (Georgeotwn University) and Dr. Vivek Bhalla (Stanford University) for helpful discussions.

