## Supplemental Figure 1, Supplemental Figure 2 for "Coordinated Regulation of Renal Glucose Reabsorption and Gluconeogenesis by mTORC2 and Potassium"

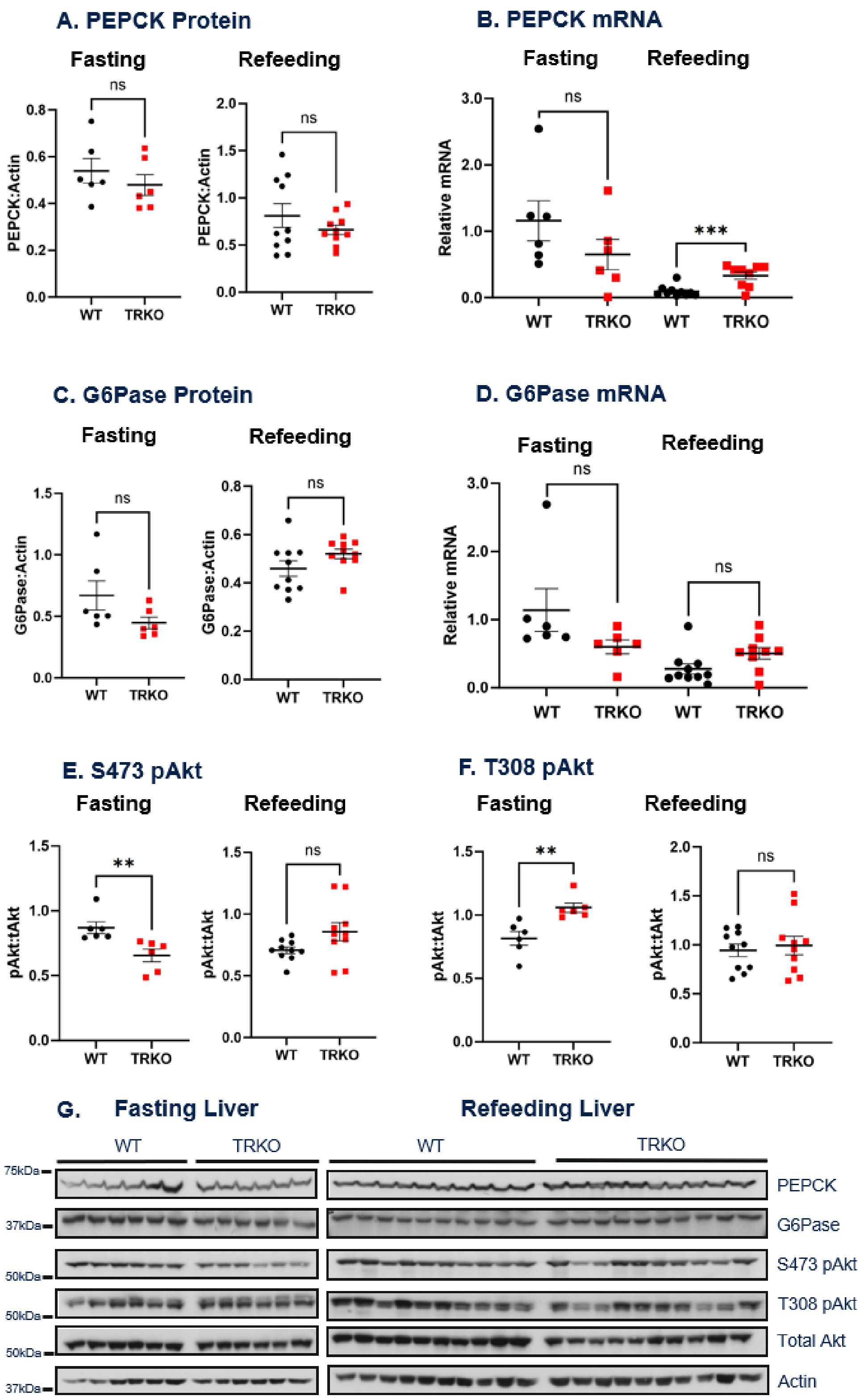


**Supplemental Figure 1: Hepatic gluconeogenic enzymes and mTORC2 signaling in TRKO and WT mice after fasting and refeeding.** (A) Western blots of whole liver homogenates for

TRKO and WT mice after 18 hours of fasting (left) and 4 hours of refeeding (right) on a normal 0.5% $K^+$ diet. (B) Quantification of PEPCK bands from Western blots and (C) relative PEPCK mRNA after fasting and refeeding in TRKO and WT mice. (D) Quantification of G6Pase bands and (E) relative G6Pase mRNA after fasting and refeeding. (F-G) Quantification of pAkt bands from Western blots after fasting and refeeding in TRKO and WT mice. PEPCK and G6Pase were normalized to actin for analysis; S473 pAkt and pT308 Akt were normalized to total Akt for analysis. Relative mRNA was calculated using the $2^{-\Delta\Delta Ct}$ method after normalizing to actin and using fasted WT mice as the reference. All values are mean ±SEM; **P <0.01, ***P<0.001, ns not significant by t-test. n=6–10 per group for all experiments.

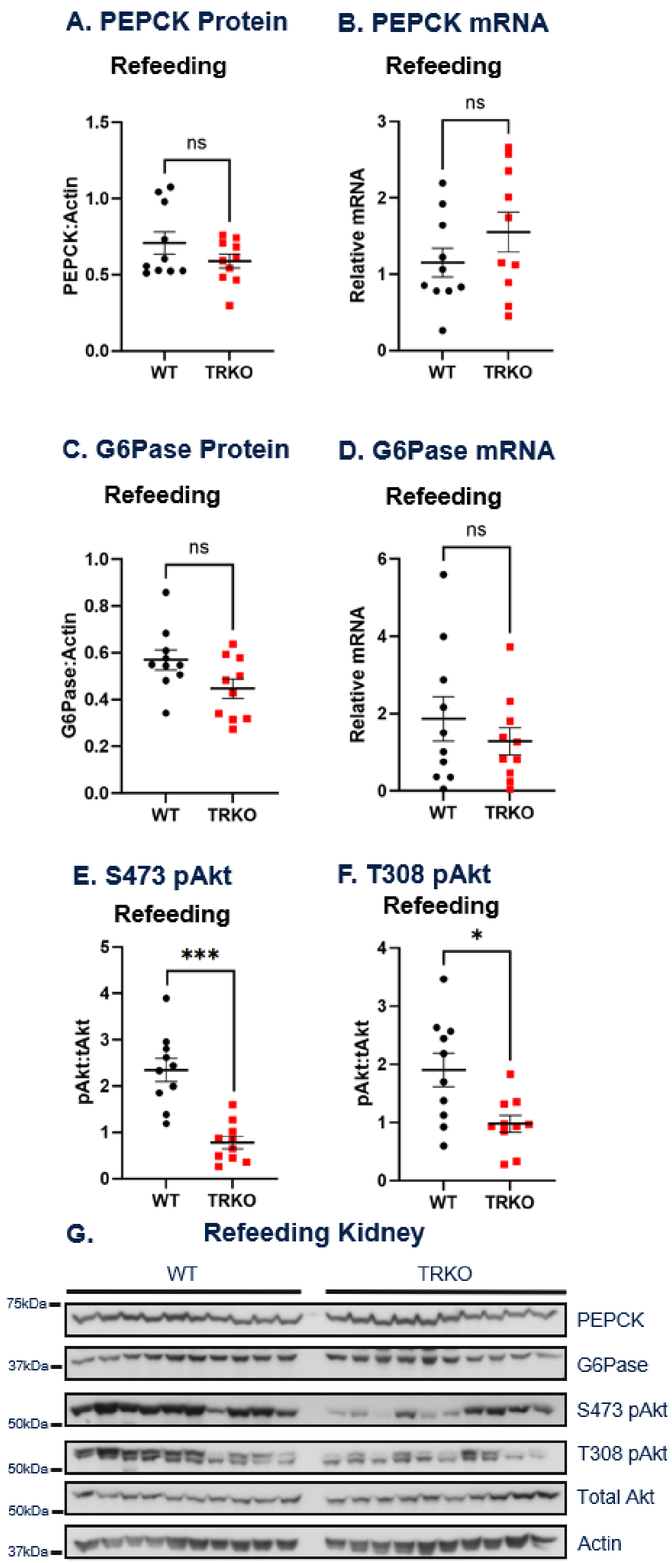


**Supplemental Figure 2: Hepatic gluconeogenic enzymes and mTORC2 signaling in TRKO and WT mice after refeeding on a high K$^+$ diet.** (A) Western blots of whole liver homogenates for TRKO and WT mice after 18 hours of fasting and 4 hours of refeeding on a high 3% $K^+$ diet. (B) Quantification of PEPCK bands from Western blots and (C) relative PEPCK mRNA after refeeding in TRKO and WT mice. (D) Quantification of G6Pase bands and (E) relative G6Pase mRNA after refeeding. (F-G) Quantification of pAkt bands from Western blots after refeeding in TRKO and WT mice. PEPCK and G6Pase were normalized to actin for analysis; S473 pAkt and pT308 Akt were normalized to total Akt for analysis. Relative mRNA was calculated using the $2^{-\Delta\Delta Ct}$ method after normalizing to actin and using fasted WT mice as the reference. All values are mean ±SEM; *P <0.05, **P <0.01, ***P<0.001, ns not significant by t-test. n=6–10 per group for all experiments.
